# Decoding of the speech envelope from EEG using the VLAAI deep neural network

**DOI:** 10.1101/2022.09.28.509945

**Authors:** Bernd Accou, Jonas Vanthornhout, Hugo Van hamme, Tom Francart

## Abstract

To investigate the processing of speech in the brain, commonly simple linear models are used to establish a relationship between brain signals and speech features. However, these linear models are ill-equipped to model a highly-dynamic, complex non-linear system like the brain, and they often require a substantial amount of subject-specific training data. This work introduces a novel speech decoder architecture: the Very Large Augmented Auditory Inference (VLAAI) network.

The VLAAI network outperformed state-of-the-art subject-independent models (median Pearson correlation of 0.19, p < 0.001), yielding an increase over the well-established linear model by 52%. Using ablation techniques we identified the relative importance of each part of the VLAAI network and found that the non-linear components and output context module influenced model performance the most (10% relative performance increase). Subsequently, the VLAAI network was evaluated on a holdout dataset of 26 subjects and publicly available unseen dataset to test generalization for unseen subjects and stimuli. No significant difference was found between the holdout subjects and the default test set, and only a small difference between the default test set and the public dataset was found. Compared to the baseline models, the VLAAI network still significantly outperformed all baseline models on the public dataset. We evaluated the effect of training set size by training the VLAAI network on data from 1 up to 80 subjects and evaluated on 26 holdout subjects, revealing a logarithmic relationship between the number of subjects in the training set and the performance on unseen subjects. Finally, the subject-independent VLAAI network was fine-tuned for 26 holdout subjects to obtain subject-specific VLAAI models. With 5 minutes of data or more, a significant performance improvement was found, up to 34% (from 0.18 to 0.25 median Pearson correlation) with regards to the subject-independent VLAAI network.

## Introduction

Recent research has focused on detecting neural tracking of speech features in EEG to understand how speech is procesed by the brain^1–3^. Neural tracking has been found for multiple acoustic representations of speech, such as the spectrogram^2, 4^ or envelope representations^1, 3, 5, 6^. Additionally, neural tracking has been shown for higher order representations such as semantic dissimilarity, cohort entropy, word surprisal, and phoneme surprisal^7–10^. Diagnostic tests can be developed that exploit the neural tracking of these features^11^. The speech envelope, for example, has been successfully linked to speech understanding^3, 6, 12^, and atypical phonological tracking has also been linked to dyslexia^13^. Neural tracking can be detected through various methodologies. In the current literature, models are used to either reconstruct the speech feature from the EEG (backward modelling)^1, 3^, predict the EEG (or a single channel of EEG) from the speech feature^14^, or transform both EEG and speech features (cf. the match-mismatch paradigm)^15, 16^.

Most commonly linear models are used^1–3, 14^. Unfortunately, the reconstruction scores are low (correlation of 0.1-0.2 between actual and reconstructed envelope for subject-specific linear decoders, 0.05 for subject-specific linear forward models) with high inter-subject variability. Subject-independent models trained on a separate dataset of other subjects would be preferable as no training data for the model has to be collected^6^, but reconstruction scores are even lower for linear models in the subject-independent setting, rendering them less useful for analysis than their subject-specific counterparts^17^.

Deep, non-linear artificial neural networks have been proposed as an alternative over linear models to model the complex non-linear brain^5, 18–21^. Recently, deep learning methods have been successfully applied to the match/mismatch paradigm^5, 20^. In this paradigm, a (non-)linear transformation of the EEG is compared to a (non-)linear transformation of a time-aligned/matched stimulus segment and a non-time-aligned/mismatched segment. The task of the model is then to identify which of the two proposed stimulus segments was time-aligned with the EEG. This method has been successfully linked to speech intelligibility^6^. Following recent advances in the match-mismatch paradigm, Thornton et al.^17^ have also shown improvements in decoding performance using neural networks in subject-specific and subject-independent settings. While deep learning is a popular method to learn complex patterns from considerable amounts of data, the low signal-to-noise ratio for auditory EEG (−10 to −20dB SNR) poses significant challenges.

We present a new decoding neural network named the *Very Large Augmented Auditory Inference* (VLAAI) network, which improves decoding performance far beyond linear methods and beyond the results of Thornton et al.^17^

## Results

The VLAAI network was compared to previously published state-of-the-art subject-independent models in subsection Comparison with baselines. Three models were chosen: a linear decoder and 2 artificial neural networks proposed by^17^: the CNN (a convolutional neural network based on EEGNET^22^) and FCNN (a multilayer perceptron based on De Taillez et al.^18^) All models try to reconstruct the stimulus speech envelope from EEG across subjects. An ablation study was performed to identify which parts of the VLAAI network are responsible for what part of the decoding performance (see subsection Ablation Study), followed by a series of experiments to test generalization (see subsection Generalization and Influence of amount of subjects/data seen during training) and subject-specific finetuning (see subsection Fine-tuning).

### Comparison with baselines

The models described in section Models are trained with the training set of the single-speaker stories dataset. The single-speaker stories dataset consists of 80 subjects listening to 1 hour and 46 minutes of continuous natural speech on average (approximately 141 hours in total, see also section Dataset). Subsequently, the models are evaluated on the test set of the single-speaker stories dataset. This test set contains the same subjects as the training set but for unseen stimuli segments. The resulting reconstruction scores were averaged across stimuli per subject. Model results were compared using a Wilcoxon signed-rank test with Holm-Bonferroni correction.

Figure 1 displays the resulting reconstruction scores. The FCNN model of Thornton et al.^17^ performed significantly worse than all other models (median Pearson r = 0.06, p<0.001). VLAAI significantly outperformed all baseline models (median Pearson r = 0.19, p<0.001), a relative improvement of 52% compared to the state-of-the-art linear model.

**Figure 1.**
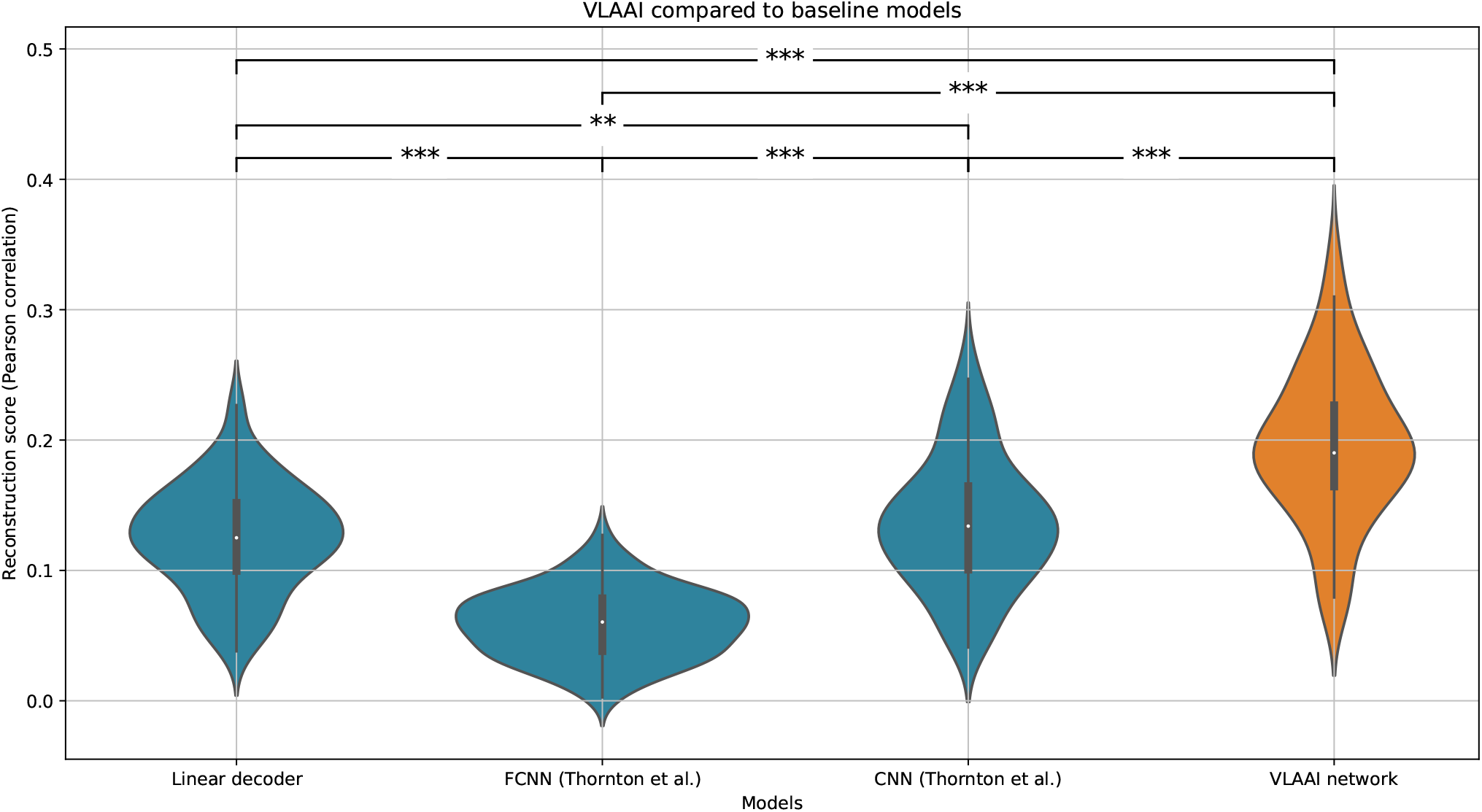
Comparison of the VLAAI network with the baseline models: a subject-independent linear model, and the FCNN and CNN models presented by^17^. Each point in the violin plot is the reconstruction score for a subject (80 subjects in total), averaged across stimuli. The FCNN performs significantly worse than the other models. No significant difference is found between the linear decoder and the CNN’s performance. The VLAAI network significantly outperforms all baseline models (*p* < 0.001), a relative improvement of 52% compared to the linear decoder. (n.s.: p ≥ 0.05, *: 0.01 ≤ p < 0.05, **: 0.001 ≤ p <0.01, ***: p < 0.001)

While the results of Thornton et al.^17^ could be replicated using their own provided data and code (https://github.com/mike-boop/mldecoders), the FCNN model performed worse when applied to our datasets. A possible explanation is that the hyperparameters found to be optimal for population models for their datasets were not as effective for our data.

### Ablation Study

It is notoriously hard to understand how non-linear artificial neural networks work. We conducted an ablation study to gain insight into what parts of the model are responsible for what part of the decoding performance.

Starting from the linear decoder baseline (cf subsection Linear decoder), more complexity is added to the model in the following steps:

1. a small (2-layer) CNN (kernel size 20 with 256 filters) with LeakyReLU^23^ activation function and layer-normalization^24^ is used, as displayed in Figure 2 (A) with *M* = 2.
2. a larger CNN is introduced, still following the structure of Figure 2 (A) but with *M* = 5, using the same CNN stack as the full VLAAI network (see also subsection Models): 256 filters for the first three convolutional layers and 128 filters for the last two convolutional layers, all with a kernel size of 8
3. The largest CNN in this step is the same as the larger CNN model of step 2, except that the filter size of all layers in the CNN stack is increased to 512.
4. The larger CNN model of step 4 is adapted to the structure displayed in Figure 2 (B), with *N* set to 4.
5. skip connections are added to the model of the fourth step to obtain the model displayed in Figure 2(C).
6. The full architecture of VLAAI is reached, as displayed in Figure 7.

**Figure 2.**
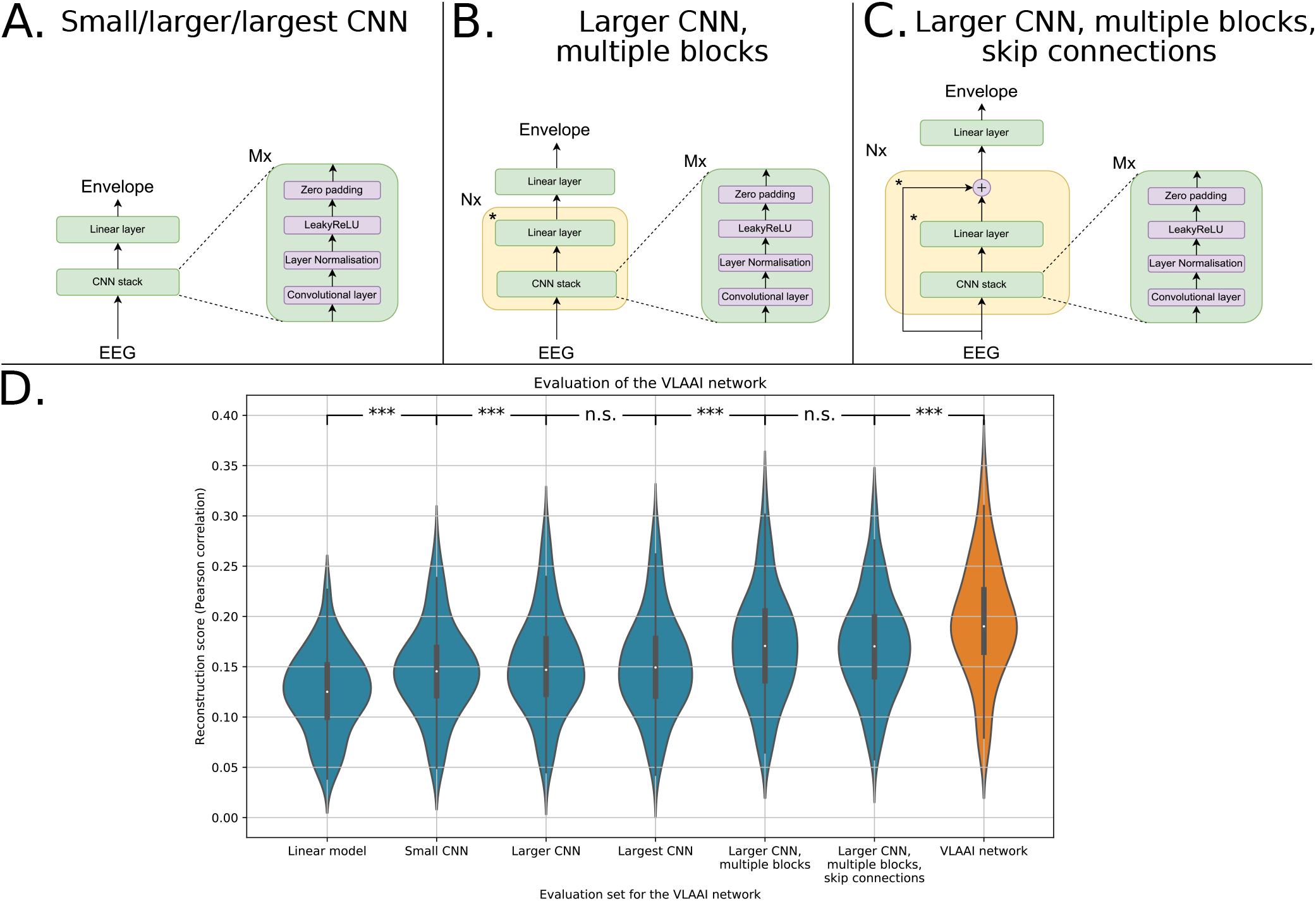
**(A)** The small/larger/largest CNN model. For the small CNN *M* = 2, the convolutional layers have a kernel size of 20 and 256 filters. The large CNN has five convolutional layers (*M* = 5) with 256 filters for the first three layers and 128 filters for the last two filters, all with a kernel size of 8. The largest CNN also has five convolutional layers (*M* = 5), but with 512 filters and a kernel size of 8. **(B)** The larger CNN, multiple blocks, following the structure of the larger CNN model. The asterisk next to the linear layer highlights that it is not present in the last repetition of that block. **(C)** The larger CNN, multiple blocks, with skip connections (step 5 in (D)). The asterisk next to the linear layer and skip connection is to highlight that it is not present in the last repetition of that block **(D)** Ablation study of the VLAAI network. Each point in the violin plot represents a reconstruction score (Pearson correlation) for a subject, averaged across stimuli. No significant difference was found between the large and largest CNN (p=0.68) and between the larger CNN with multiple blocks and the larger CNN with multiple blocks with skip connections (p=0.99). The biggest increases in reconstruction score are between the linear model and the small CNN (14% increase in median reconstruction score), the larger CNN with *N* = 1 and *N* = 4 (10% increase in median reconstruction score) and when adding the output context layer to the penultimate model to obtain the VLAAI network (10% increase in median reconstruction score).(n.s.: p ≤ 0.05, *: 0.01 ≤ p < 0.05, **: 0.001 ≤ p < 0.01, ***: p < 0.001)

**Figure 3.**
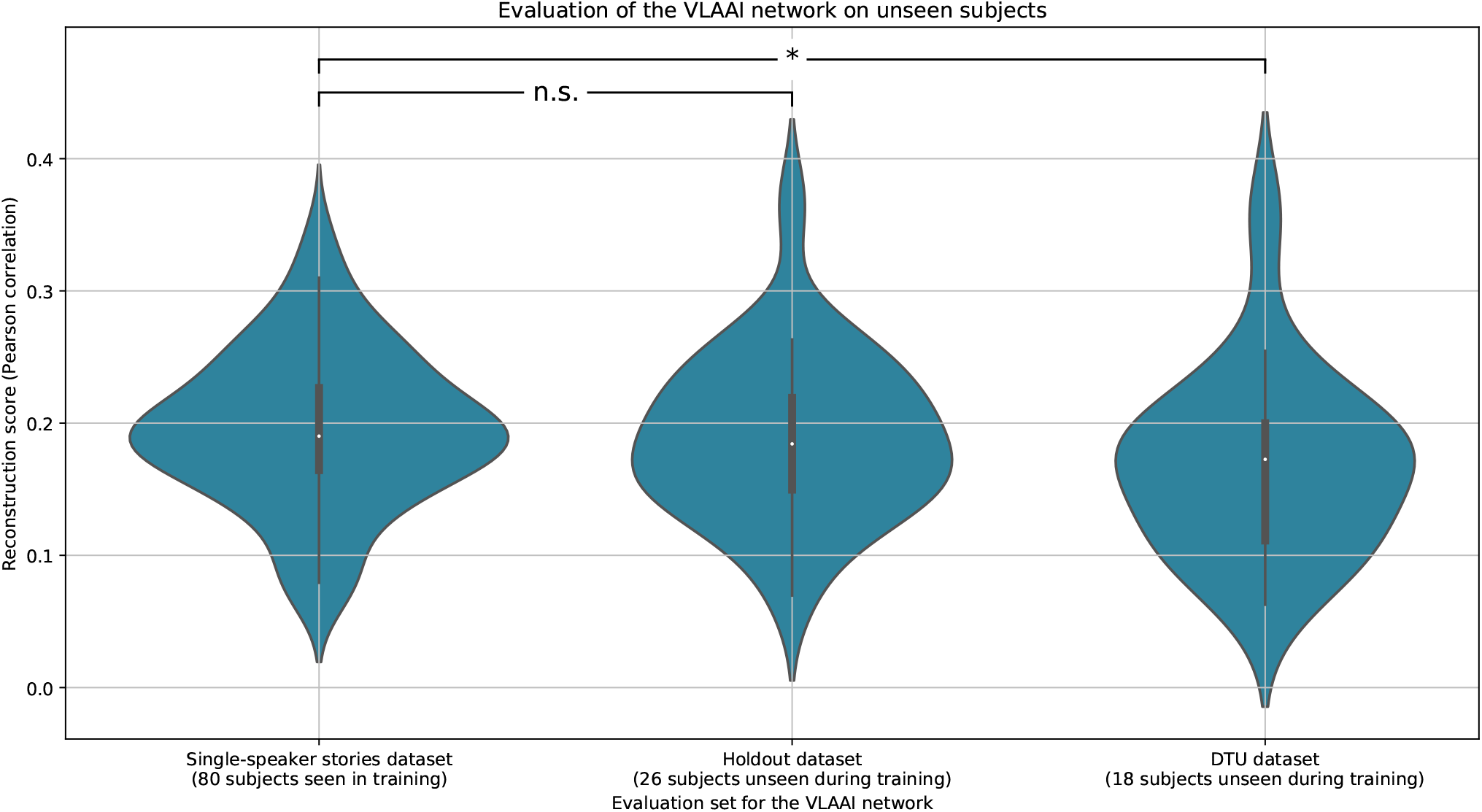
Generalization between the single-speaker stories dataset, the holdout dataset and the single-speaker trials of the publicly available DTU dataset. Each point in the violin plot is the reconstruction score for a subject, averaged across stimuli. No significant difference in reconstruction score was found between the single-speaker stories and the holdout dataset (p=0.23). A significant difference in reconstruction score was found between the single-speaker stories dataset and the DTU dataset (p=0.04), possibly due to differences in measuring protocol and/or stimulus characteristics. (n.s.: p ≤ 0.05, *: 0.01 ≤ p < 0.05, **: 0.001 ≤ p < 0.01, ***: p < 0.001)

**Figure 4.**
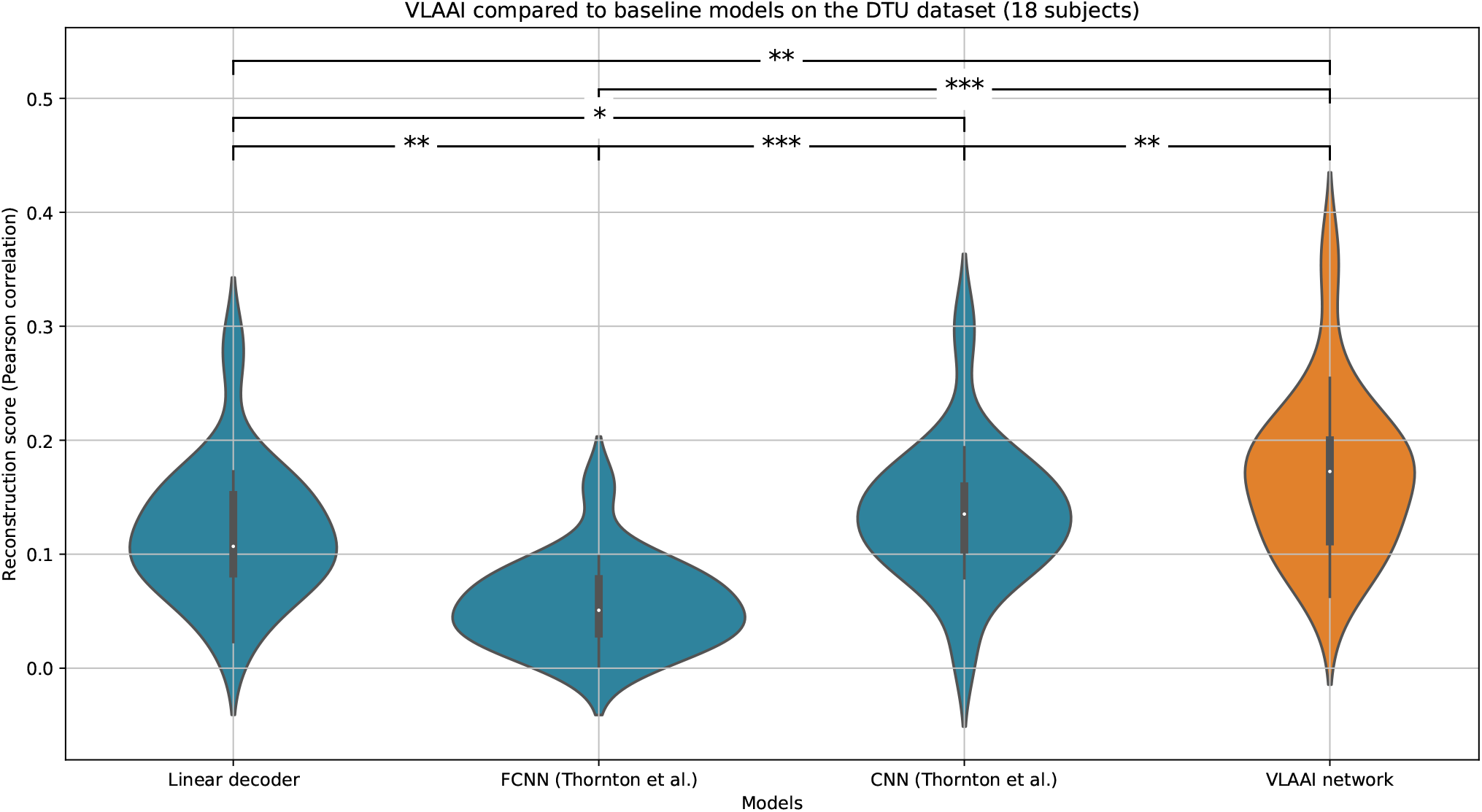
Evaluation of the baseline models and the VLAAI network on the single-speaker trials (50 seconds per trial) of the DTU dataset. While the reconstruction scores are lower for all models compared to the single-speaker stories dataset (see also Figure 1), the general conclusions still hold. The VLAAI network significantly outperforms all other baseline models (p<0.01), with a relative performance increase of 62% over the linear decoder. (n.s.: p ≤ 0.05, *: 0.01 ≤ p < 0.05, **: 0.001 ≤ p < 0.01, ***: p < 0.001)

**Figure 5.**
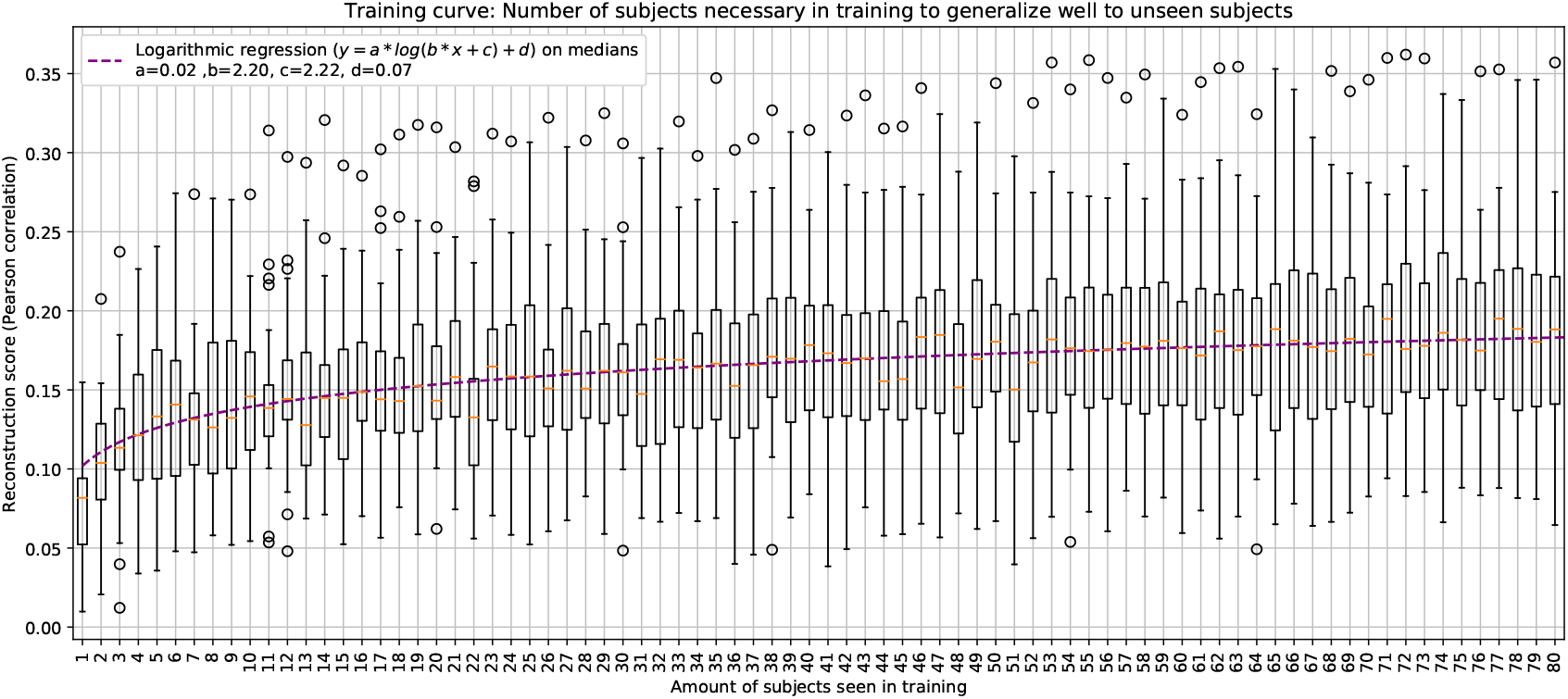
The VLAAI network was trained using 1-80 subjects of the single-speaker stories dataset and evaluated on the holdout dataset (26 subjects). Each point in the boxplot is the reconstruction score for a subject, averaged across stimuli. Median reconstruction scores increase logarithmically with the number of subjects seen during training.

**Figure 6.**
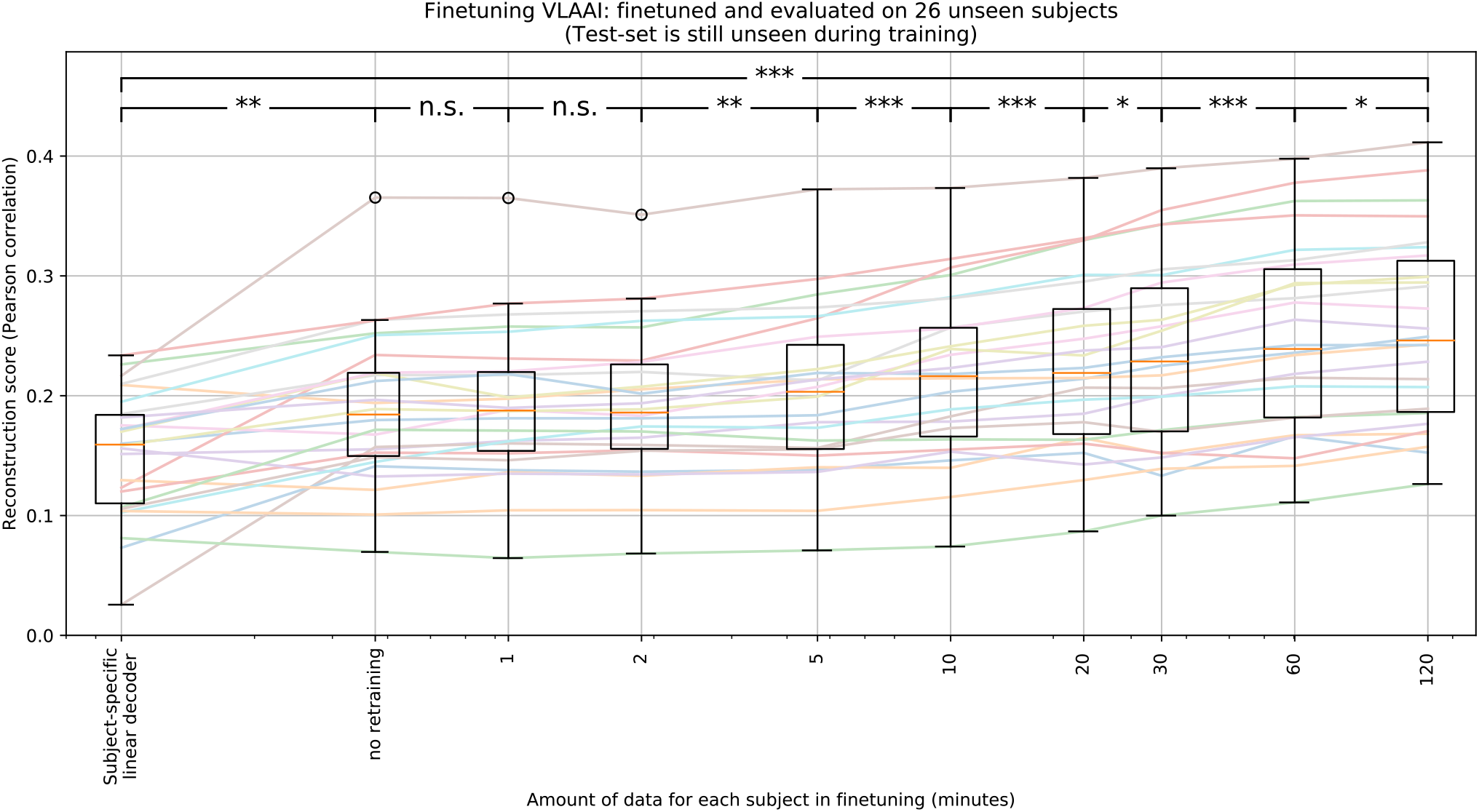
Subject-independent VLAAI, trained on the single-speaker stories dataset, fine-tuned on the subjects of the holdout dataset. Each point in the boxplot is the reconstruction score for a subject, averaged across stimuli. No significant increase is found between the subject-independent model (no fine-tuning) and models fine-tuned with 1 and 2 minutes of data. Starting with 5 minutes of available fine-tuning data, the median reconstruction score seems to increase logarithmically from 0.19 with the amount of training data to 0.25 Pearson r for 120 minutes. (n.s.: p ≥ 0.05, *: 0.01 ≤ p < 0.05, **: 0.001 ≤ p < 0.01, ***: p < 0.001)

**Figure 7.**
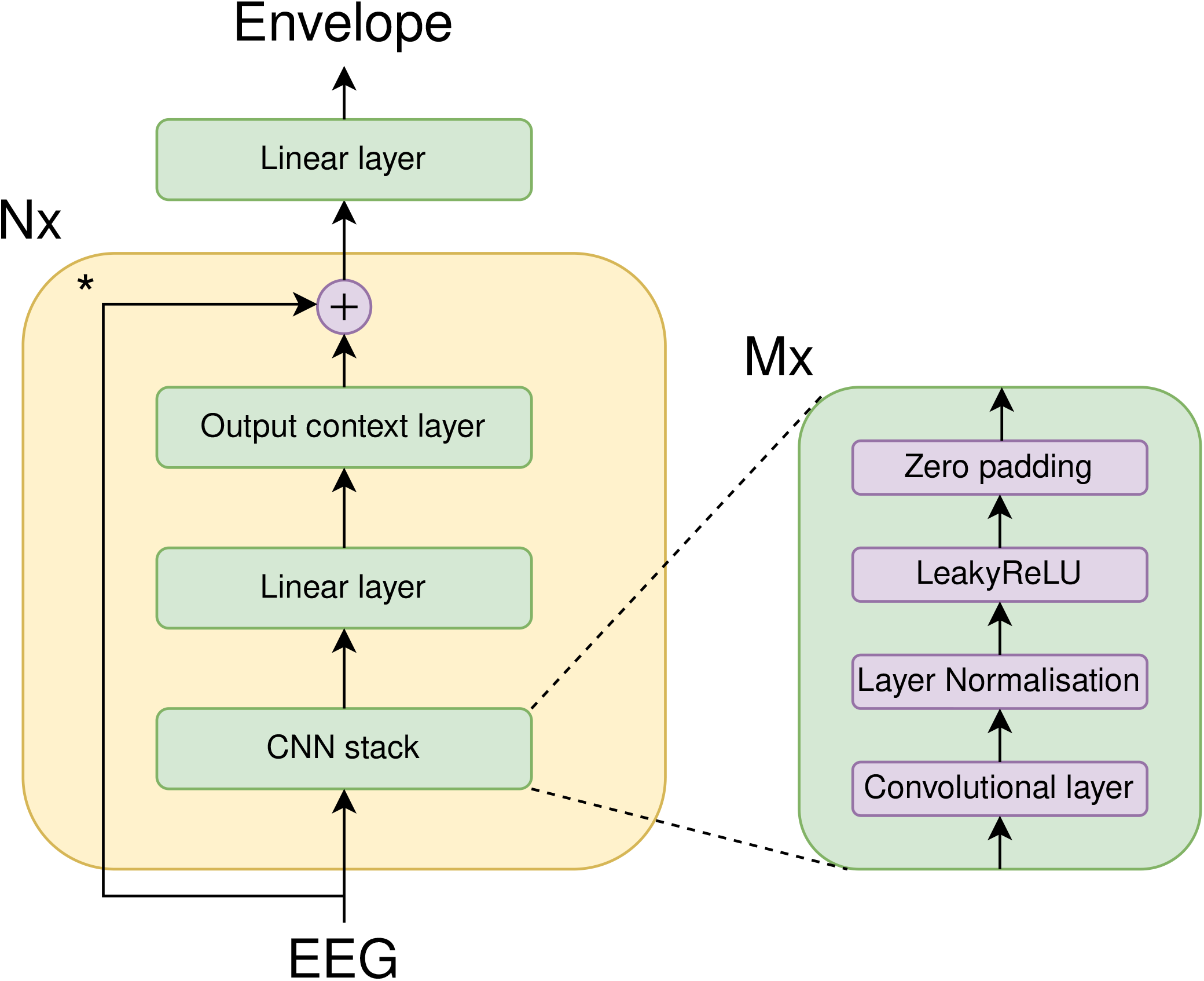
Structure of the proposed VLAAI network. The asterisk next to the skip connection indicates that it is not present in the last repetition of that block.

Models in subsequent steps were compared using a Wilcoxon signed-rank test with Holm-Bonferroni correction.

As shown Figure 2(D), adding more model complexity, weights and non-linearities seem to improve performance up until step 3 (larger CNN), after which increasing the filter sizes of the model has no significant effect (p=0.61). Increasing the number of blocks from *N* = 1 to *N* = 4 delivers a big performance increase (≈a 10% increase in median reconstruction score compared to step 3). Finally, the output context layer substantially improves the performance (≈a 10% increase in median reconstruction score compared to step 5).

While adding complexity is certainly beneficial to decoding performance when the models are small, it seems that at a certain size, a point of diminishing returns is reached with regard to adding weights. The increased performance added by the output context layer suggests that the model can extract beneficial information based on the previous output context.

### Generalization

Subject-independent models should be able to generalize well across different subjects, even if they are unseen in training. This generalization is crucial, as testing time is expensive.

To evaluate the across-subject generalization, the VLAAI network is trained on 80 subjects of the single-speaker stories dataset and evaluated on the test set of these 80 subjects, compared to the test set of the 26 subjects in the holdout dataset, as well as the single-speaker trials (50 seconds per trial) of the 18 subjects of the publicly available DTU dataset^25^. The results on the test-set of the single-speaker stories dataset are compared to the test-set of the holdout and DTU dataset.

In Figure 3, the decoding performance of the VLAAI network on the test set of the 80 subjects seen during training and the test set of the 26 subjects of the holdout dataset (unseen during training) is shown. Reconstruction scores were not significantly different (p=0.23, 95% confidence interval of the difference=[-0.02, 0.04]) for the test set of the singe-speaker stories dataset compared to the test set of the holdout dataset. There was a significant difference between performance on the test set of the single-speaker stories dataset and the DTU dataset (decrease from 0.19 median Pearson r to 0.17 Pearson r, p=0.04). Possible causes for this difference could be the difference in stimulus characteristics (e.g. intonation, language,..), measurement protocol or systematic differences in EEG cap placement. Additionally, to evaluate if the findings of our first experiment still hold, the baseline models (linear decoder, CNN and FCNN) of the first experiment (subsection Comparison with baselines) and the VLAAI network are evaluated on the DTU dataset (see Figure 4). While reconstruction scores decrease for all models compared to subsection Comparison with baselines, the general conclusions still hold. The VLAAI network significantly outperforms all other models (p<0.01), with a relative performance increase of 61% over the linear decoder.

### Influence of amount of subjects/data seen during training

Subject-independent models must learn to extract subject-independent patterns and characteristics of neural tracking of the stimulus. This requires a training set with many subjects to prevent overfitting on specific characteristics of the subjects seen in training.

To assess the influence of adding data of new subjects to the training set, the VLAAI network was trained on 1-80 subjects of the single-speaker stories dataset and subsequently evaluated on the test-set of the 26 subjects of the holdout dataset. A logarithmic function (*Pearsoncorrelation = a* * *log*(*b* * (*amount of subjects in training*)+*c*)+*d*) is fitted on the median decoding performance of all models to characterise the relationship between decoding performance and the number of subjects used in training, using *scipy.optimize.curve_fit*^26^.

The results are visualized in Figure 5, the median correlation increases logarithmically with the number of subjects seen in training (80). The most dramatic increase (the 90th percentile of the increase) is seen for the first 9 subjects (from 0.08 Pearson r to 0.14 Pearson r, a relative increase of 85%). This logarithmic relationship between the number of subjects and decoding performance can be used to extrapolate and estimate what decoding performance can be reached by including more subjects in training (e.g, approximately 20000 subjects would yield a median Pearson correlation of 0.30). Nevertheless, a plateau in decoding performance is expected to be reached when the models’ weights are saturated. The found relationship could be different for other, less homogeneous datasets.

### Fine-tuning

In current literature, most of the focus has been on subject-specific models, as they are easy to train and well suited for diagnostic applications. Subject-independent models can, however, be fine-tuned on a specific subject after subject-independent training. Starting from a previously trained subject-independent decoder can increase decoding performance as the already found general patterns are adapted to be optimal for a single subject.

The VLAAI network was trained on the training set of the single-speaker stories dataset and subsequently fine-tuned separately on the training set of the subjects of the holdout dataset for different amounts of training data per subject (i.e. 1 minute, 2 minutes, 5 minutes, 10 minutes, 20 minutes, 30 minutes, 1 hour and 2 hours), taken uniformly from the available recordings for each subject. To prevent overfitting, the batch size was reduced to 1, and the learning rate was lowered to 10^-4^. The test set of the holdout dataset was used for evaluation. To compare these fine-tuned subject-specific VLAAI models to the state-of-the-art for subject-specific decoding, subject-specific linear decoders were trained (with ridge regression using an integration window of 250 ms) on the training set of the holdout dataset and evaluated on the test-set of the holdout dataset. The performance of subsequent models as a function of the amount of fine-tuning data was compared using a Wilcoxon signed-rank test with Holm-Bonferroni correction.

As seen in Figure 6, even without fine-tuning, the subject-independent VLAAI model already significantly outperforms the subject-specific linear decoders (median Pearson r of 0.18 vs 0.16 respectively, p<0.01). No significant increase is found from fine-tuning (the subject-independent model) with up to 2 minutes of data. The reconstruction scores of all other models were significantly different for subsequent amounts of training data (p<0.01). From 5 minutes onwards, the performance seems to increase logarithmically, reaching a top median reconstruction score of 0.25 Pearson r, yielding an even higher relative performance increase of 55% over the subject-specific linear decoders (p<0.001).

This implies that the VLAAI network can derive subject-specific patterns better suited for decoding than the already extracted subject-independent patterns, setting new state-of-the-art reconstruction scores for subject-dependent models.

## Discussion

This study introduced the new VLAAI network, compared it to baseline models from^17^ and evaluated subject generalization and subject-specific fine-tuning. In subsection Comparison with baselines, the VLAAI network was shown to significantly outperform the proposed baseline models (see also subsection Models). As shown by^17^, increasing the number of weights does not necessarily increase decoding performance. In section Ablation Study, diminished returns are indeed achieved in decoding performance when using standard non-linear artificial neural network architectures: the biggest increases in decoding performance of the VLAAI network are due to the use of a bigger model with non-linearities, the stacking of multiple blocks and the output context layer. The non-linearities might help in modelling the highly complex and non-linear auditory processing in the brain, while the output context layer can use the previous output context to refine predictions (i.e. some predictions might be implausible when taking the previous context into account). The benefit of taking previous context into account was already suggested by^18^. In contrast to^18^, the output context layer is built into the model and can therefore also be leveraged during evaluation. The higher decoding performance of VLAAI can be used to reveal the effects of auditory processing in EEG that were previously hidden. A downside of the VLAAI network, is that the model itself cannot be easily interpreted, in contrast to forward linear models that can be interpreted as temporal response functions^1, 27^. However, the same is true for backward linear models, which are frequently used^28^.

In subsection Generalization, no significant decline in reconstruction scores was found for subjects unseen during training, but a small decrease in median Pearson r was found for the DTU dataset (from 0.19 Pearson r to 0.17, p=0.04). While the DTU dataset was recorded using a similar EEG system (BioSemi ActiveTwo), the measurement paradigm and stimulus characteristics differed from the single-speaker stories dataset (i.e. Shorter trials were presented and were part of a bigger auditory attention detection paradigm). Differences in auditory processing for different languages might also have influenced the results, as their stimuli were narrated in Danish and presented to native Danish listeners, while the single-speaker stories dataset contained Dutch stimuli presented to native Dutch speakers. In addition, systematic difference in EEG cap placement could have influenced the results. Further research is needed to track down the exact reason for this small difference in reconstruction performance. Nevertheless, correlation scores of the VLAAI network are still high compared to the scores of the baseline models when evaluated on the DTU dataset (p<0.01), setting a new state-of-the-art for subject-independent models.

Fine-tuning the VLAAI network shows that it can be transformed effectively from a subject-independent to a subject-specific model with as little as 5 minutes of additional data per subject. While the subject-independent model already significantly outperformed the subject-specific linear decoders (0.19 median Pearson r compared to 0.16 median Pearson r respectively, p<0.01), fine-tuning can increase the performance to even higher levels (median Pearson r of 0.25). This increased performance might allow uncovering previously undetectable neural processes and improve measurement efficiency, showing promise for the VLAAI network to be used in applications such as diagnostic hearing tests. In future research, better results might be obtained with even less additional data by only retraining a certain subset of layers.

Decoding other speech features might give more insight into more complex stages of the auditory system, such as the neural tracking of phonemes or semantics. However, validation on datasets of different populations (people with varying levels/causes of hearing impairment, etc.), measuring paradigms and more diverse stimuli (spontaneous speech, etc.) are still required to move towards a robust clinical application.

In conclusion, this paper proposes a new non-linear neural network architecture, which sets a new state-of-the-art in decoding the speech envelope from EEG in both a subject-independent setting (median reconstruction scores of 0.19 Pearson r, a relative improvement of 52% over the subject-independent linear decoder), and a subject-specific setting (median reconstruction scores of 0.25 Pearson r after fine-tuning, a relative improvement of 54% over the state-of-the-art subject-specific linear decoder models).

Our code and pre-trained models are available on https://github.com/exporl/vlaai, allowing other researchers to easily use the VLAAI network for their research/experiments.

## Materials and methods

### Models

The new VLAAI network was compared to three baseline models: a linear decoder, the FCNN network and the CNN network from^17^. Each model was trained for at most 1000 epochs, with early stopping using a patience factor of 5 and a minimum delta of 10^-4^.

#### Linear decoder

The linear decoder reconstructs the speech envelope from EEG by using a linear transformation across all channels and a certain time/integration window. Contrary to most studies, the linear decoder here is trained subject-independently with negative Pearson r as a loss function to have a fair comparison with the other proposed subject-independent models. In preliminary experiments, Pearson r yielded better decoding performance than MSE in the subject-independent scenario. An integration window of 500 ms was used in all experiments. The linear decoder was trained using Adam^29^, using a learning rate of 10^-3^ on overlapping windows of 5 seconds (80% overlap) and a batch size of 64. The linear decoder was implemented in Tensorflow version 2.3.0^30^.

For the ablation experiment (subsection Ablation Study), subject-specific linear decoders were used. These linear decoders were trained in a similar fashion as^1, 3^: using ridge regression with laplacian regularisation^31^ and an integration window of 250 ms. This was implemented using the *mne.decoding.ReceptiveField* class from MNE^32^. 15 ridge parameters were sampled logarithically from 10^-7^ to 10^7^, and the model performing best on the validation set was chosen for evaluation.

#### FCNN

The second proposed baseline is the FCNN model introduced in Thornton et al.^17^, using the found optimal hyperparameters for the population model reported in their work. This model is a multilayer perceptron, with weight decay applied to the hidden layers and tanh non-linearities, batch normalization^33^ and dropout^34^ applied subsequently. The number of units of each layer was 2363, 1576 and 788, respectively. The drop rate for each dropout layer was fixed at 45%. The model was trained using segments of 400 ms without overlap. The FCNN was trained with Nadam^35^ using a learning rate of 2^-3^, using negative Pearson r as a loss function. The batch size used during training was 256. The model, training and evaluation code for the FCNN models were used and adapted from the author’s GitHub (https://github.com/mike-boop/mldecoders) using Pytorch 1.10.0^36^.

#### CNN

The final baseline model is the CNN model introduced in Thornton et al.^17^, using the found optimal parameters for the population model. This model is based on the EEGNET architecture^22^. The model consists of 4 convolutional layers: a temporal convolution and a depthwise convolution^37^, which combines the channels of the temporal convolution, followed by another depthwise convolution across the time dimension. After each depthwise convolution, batch normalisation^33^, Exponential linear units (ELU) non-linearities^38^, average pooling (with a factor of 5 and 2 respectively) and spatial dropout (with a drop rate of 20%)^34^ were applied. Finally, the output of the last convolution is flattened, and all samples are combined with a fully connected layer with a linear activation. The CNN was trained with Nadam^35^ using a learning rate of 2^-3^, with negative Pearson r as a loss function. The batch size used during training was 256. As with the FCNN model, code for the CNN model architecture, training and evaluation was adapted from the author’s GitHub (https://github.com/mike-boop/mldecoders).

#### VLAAI

We propose a new model, called the VLAAI network, which consists of multiple (*N*) blocks, consisting of 3 different parts (see also Figure 7). The first part is the CNN stack, a convolutional neural network. This convolutional neural network consists of *M*=4 convolutional layers. The three first layers have 256 filters, while the last two layers have 128 filters. Layer normalization^24^, a LeakyReLU^23^ activation function, and zero-padding with 7 samples at the end of the sequence are applied after every layer. The second part is a simple, fully connected layer of 64 units, which recombines the output filters of the CNN stack. The last part is the output context layer. This special layer enhances the predictions made by the model up until that point by taking the previous context into account and combining it with the current sample. Concretely, this block makes a combination in the time dimension of the previous 31 samples, combined with the current sample and predict a better version of the current sample. A convolutional layer with a kernel of 32 and 64 filters is used, combined with a LeakyReLU non-linearity and layer normalization. Taking the output context into account was also advantageous during training in De Taillez et al.^18^, although the implementation in VLAAI is integrated into the model and used in both training and evaluation. At the end of each block except the last, a skip connection is present with the original EEG input. After the last block, the linear layer at the top of the VLAAI model combines the 64 filters of the output context layer into a single speech envelope. To prevent overfitting, weights from the CNN stack and autoregression layer are shared across blocks. The VLAAI network was trained with Adam using a learning rate of 10^-3^, with negative Pearson r as a loss function. The batch size used during training was 64.

Code for the VLAAI network and pre-trained models can be found at https://github.com/exporl/vlaai.

#### Dataset

Our own dataset (the single-speaker stories dataset) is used to evaluate the VLAAI network and the baseline models. A subset of the publicly available DTU dataset is also used to extensively evaluate the generalizability of VLAAI and the baseline models.

The single-speaker stories dataset contains 106 normal-hearing participants between 18 and 30 years old. Participants signed informed consent for this study, approved by the Medical Ethics Committee UZ KU Leuven/Research (KU Leuven, Belgium) with reference S57102. Firstly, participants were asked to fill in a questionnaire, confirming that they have no neurological or auditory conditions. Secondly, the participants’ hearing was tested using a pure-tone audiogram and a Flemish MATRIX test. Participants with hearing thresholds of >30dBHL were excluded.

Following the screening procedure, the EEG of participants was measured while they listened to 2-8 (on average 6) single-speaker stories. Longer stories were partitioned into multiple parts. Each part was approximately 15 minutes long. 2 out of the 10 parts were presented with speech-weighted noise at 4dB SNR, but were excluded from this study. Participants were notified before listening that they had to answer a question about the content of the story after listening as an incentive to pay close attention to the contents of the story. Throughout the recording session, participants were given short breaks.

A subset of this dataset was also used in Accou et al.^5, 6^, Monesi et al.^4, 20^ and Bollens et al.^21^ This dataset contains approximately 188 hours of EEG recordings (on average 1 hour and 46 minutes per subject) in total.

Data from 26 (randomly chosen) subjects was designated as holdout data (the holdout dataset), while data from the remaining 80 subjects were used as standard training-, validation- and test-set (the single-speaker stories dataset). The holdout dataset contains 46 hours of EEG recordings, while the single-speaker stories dataset contains 142 hours of EEG data (1 hour and 46 minutes of speech on average for both datasets).

EEG data were collected at a sampling rate of 8192 Hz using a BioSemi ActiveTwo setup (Amsterdam, Netherlands). Electromagnetically shielded ER3A insert phones, an RME Multiface II sound card (Haimhausen, Germany), and a computer running APEX^39^ were used for stimulation. The stimulation intensity of all stimuli was fixed at 62 dBA for each ear. All recordings were performed in a soundproofed and electromagnetically shielded booth.

The DTU dataset^25^ contains EEG recordings of 18 Danish subjects that listened to natural speech in Danish spoken by 1 or 2 speakers in different reverberation settings. This dataset was also used by^40, 41^. For our study, we used only the single-speaker trials. Each trial is approximately 50 seconds long, resulting in 500 seconds of data per subject. This data is only used for evaluation, not for training.

### Preprocessing

EEG data was high-pass filtered with a 1st order Butterworth filter with a cut-off frequency of 0.5Hz using zero-phase filtering by filtering the data in both the forward and backward direction. The speech envelope was estimated using a gammatone filterbank^42, 43^ with 28 filters spaced by equivalent rectangular bandwidth with center frequencies of 50 Hz to 5 kHz. Subsequently, the absolute value of each sample in the filters was taken, followed by exponentiation with 0.6. Finally, the mean of all filters was calculated to obtain the speech stimulus envelope^44^. After downsampling EEG and speech envelopes to 1024 Hz, eyeblink artefact rejection was applied to the EEG using a multi-channel Wiener filter^45^. Next, the EEG was re-referenced to a common average. Finally, both EEG and speech envelopes were downsampled to 64 Hz.

Each EEG recording was split into a training, validation and test set, containing 80%, 10% and 10% of the recording, respectively. The validation and test set were extracted from the middle of the recording to avoid artefacts at the beginning and end of the recording. The mean and variance of each channel of EEG and the speech envelope were calculated on the training set. The EEG and envelope were then normalized by subtracting the mean from each channel and dividing by the variance for the training, validation and test set. As the DTU dataset is only used for evaluation, each trial is normalized separately and used as the test set.

All preprocessing steps were implemented in Matlab 2021a (Natick, USA), except the splitting and normalization, which were done in Python 3.6 using Numpy^46^.

In all experiments, training and testing were performed on 5-second windows with 80% overlap unless specifically stated otherwise. Reported p-values for tests using Holm-Bonferroni correction for multiple comparisons are corrected p-values.

## Data availability statement

The single-speaker stories dataset analyzed during the current study is available from the corresponding author on reasonable request and in compliance with the participant’s consent. The DTU dataset analyzed during the current study is available in the Zenodo repository,https://doi.org/10.5281/zenodo.1199011.

## Additional information

### Competing interests

The authors declare no competing interests.

## Acknowledgements

The authors thank Amelie Algoet, Jolien Smeulders, Lore Kerkhofs, Sara Peeters, Merel Dillen, Ilham Gamgami, Amber Verhoeven, Lies Bollens and Wendy Verheijen for their help with data collection. Special thanks to Simon Geirnaert and Tom Francart for their help naming the model.

The research conducted in this paper is funded by KU Leuven Special Research Fund C24/18/099 (C2 project to Tom Francart and Hugo Van hamme), by a PhD grant (1S89620N) of the Research Foundation Flanders (FWO) and from the European Research Council (ERC) under the European Union’s Horizon 2020 research and innovation program (grant agreement No 637424, ERC Starting Grant to Tom Francart)

## Author contributions statement

Bernd Accou conceived the network architecture, ran the experiments and analyzed the results. Jonas Vanthornhout, Hugo Van hame and Tom Francart helped design experiments and gave feedback on the analysis of the results. All authors reviewed and contributed to the manuscript.

## Notes

https://github.com/exporl/vlaai

### Competing Interest Statement

The authors have declared no competing interest.

### Summary of Updates

Small changes + submission to Scientific Reports

